# Mechanism-selective deep mutational scanning distinguishes *ERCC2* disease phenotypes

**DOI:** 10.64898/2026.09.24.754105

**Authors:** Hasan Çubuk, Vahid Aslanzadeh, Yifei Shang, Marcin Plech, Ankit Pathak, Grzegorz Kudla, Joseph A. Marsh

## Abstract

Pathogenic *ERCC2* variants cause xeroderma pigmentosum (XP), trichothiodystrophy (TTD) or both, yet variant effect scores are usually interpreted only as measures of pathogenicity rather than of which disease mechanism is disrupted. XPD, the *ERCC2*-encoded TFIIH subunit, functions in both nucleotide excision repair and transcription. Using yeast complementation deep mutational scanning, we measured the effects of nearly all XPD amino acid substitutions. The assay was mechanism-selective: it preferentially reported transcription-associated function, with pronounced intolerance at the p44 interface, whereas many substitutions affecting DNA binding and helicase activity retained near-wild-type fitness. Accordingly, TTD variants had much lower fitness than XP variants. Computational predictors discriminated pathogenic from benign variants similarly across phenotypes, but the DMS distinguished XP from TTD variants better than all 73 predictors tested. Phenotype-specific ACMG/AMP calibration provided evidence in both directions for TTD but mainly pathogenic evidence for XP. Thus, the selectivity of functional assays, often viewed as a limitation, can reveal disease mechanisms and support phenotype-aware variant interpretation.

## Introduction

Human XPD, encoded by *ERCC2*, is a core subunit of TFIIH, a multiprotein complex that is essential for transcription initiation and nucleotide excision repair (NER). Within TFIIH, XPD undergoes dynamic structural rearrangements and changes in subunit interactions that switch it between a non-catalytic, scaffolding role during transcription and a catalytically active helicase state during NER. XPD contains two RecA-like motor domains, RecA1 and RecA2, together with an FeS domain and an Arch domain that form a channel around DNA and support helicase activity.

During NER, XPD functions as a 5′-3′ DNA helicase, with the FeS and motor domains contributing to DNA binding, ATPase activity, and strand separation. In transcription initiation, by contrast, XPD is thought to act primarily as a structural scaffold within TFIIH, linking the CDK-activating kinase (CAK) complex to the preinitiation complex (PIC) through interactions involving the Arch and RecA2 regions. XPD also contributes to cell cycle regulation and development through modulation of CDK7 activity^1–5^.

Pathogenic *ERCC2* variants are primarily associated with two distinct recessive disorders: xeroderma pigmentosum (XP) and trichothiodystrophy (TTD). These phenotypes reflect the separable roles of XPD in nucleotide excision repair (NER) and transcription. Variants that primarily impair NER cause XP, characterised by UV sensitivity and a markedly increased risk of skin cancer, whereas variants more closely associated with defects in basal transcription and TFIIH stability cause TTD, characterised by progressive neurodegeneration, recurrent infections and, in severe cases, early mortality^6–8^. However, this distinction is not absolute: several *ERCC2* variants have been associated with both XP and TTD phenotypes, and clinical presentation may depend on the biallelic combination of alleles and their residual activities. Structural and biochemical studies have further shown that disease-associated *ERCC2* variants can differentially affect helicase activity, DNA binding and interactions within TFIIH, providing a mechanistic basis for these phenotype-associated differences^2,5,9^.

Multiplexed assays of variant effect (MAVEs) measure the functional effects of many variants in parallel^10^. Deep mutational scanning (DMS) is a form of MAVE in which large numbers of amino acid substitutions are systematically assayed to generate quantitative maps of protein function^11^. MAVEs are increasingly used as functional evidence in clinical variant classification, helping to resolve variants of uncertain significance (VUS)^12^. Unlike computational variant effect predictors (VEPs), however, which aim to capture deleteriousness in general, a MAVE measures a specific molecular or cellular readout. For a multifunctional protein, that readout may depend on only a subset of its activities, so variants that disrupt other functions can appear wild-type-like. This specificity is often regarded as a limitation. However, when the functions captured by the readout correspond to a particular disease mechanism, the same property could instead allow an assay to distinguish variants according to the mechanism they disrupt; we refer to such an assay as mechanism-selective. XPD offers a natural test of this idea. In yeast complementation assays, variants of a human protein are tested for their ability to replace its yeast orthologue, so growth reports only the functions required for complementation. This approach has been used to study human disease variants in yeast^13–16^, and is particularly well suited to XPD because human *ERCC2* can functionally complement yeast *RAD3*^17^. Because NER is dispensable for yeast growth in the absence of exogenous DNA damage, a complementation assay would be expected to report transcription-associated XPD function selectively.

We therefore used yeast complementation DMS to map the effects of *ERCC2* missense variation, to test whether this assay is mechanism-selective and whether that selectivity distinguishes XP-from TTD-associated variants. We show that growth-based complementation selectively reports transcription-associated defects caused by disruption of TFIIH stability and XPD interactions with p44 and MAT1, whereas many variants affecting DNA binding, ATPase activity and NER-specific functions retain near wild-type growth. By comparing the DMS map with biochemical data, structural features and computational VEPs, we identify systematic patterns of agreement and disagreement that reveal the molecular mechanisms captured by the assay. Finally, we calibrate the DMS scores within the ACMG/AMP framework to provide phenotype-aware evidence for *ERCC2* variant interpretation, illustrating how mechanism-selective functional maps can complement computational prediction and support clinical classification.

## Results

### A yeast complementation screen maps the functional effects of ERCC2 missense variants

To systematically measure the functional effects of *ERCC2* missense variants, we expressed human XPD in a TetO7-*RAD3* yeast strain in which endogenous *RAD3* expression can be repressed by doxycycline treatment. Because *RAD3* is essential, growth under repressed conditions provides a complementation readout: *ERCC2* variants that preserve the *RAD3*-complementing function support growth, whereas damaging variants reduce growth (Fig. 1a).

**Figure 1:**
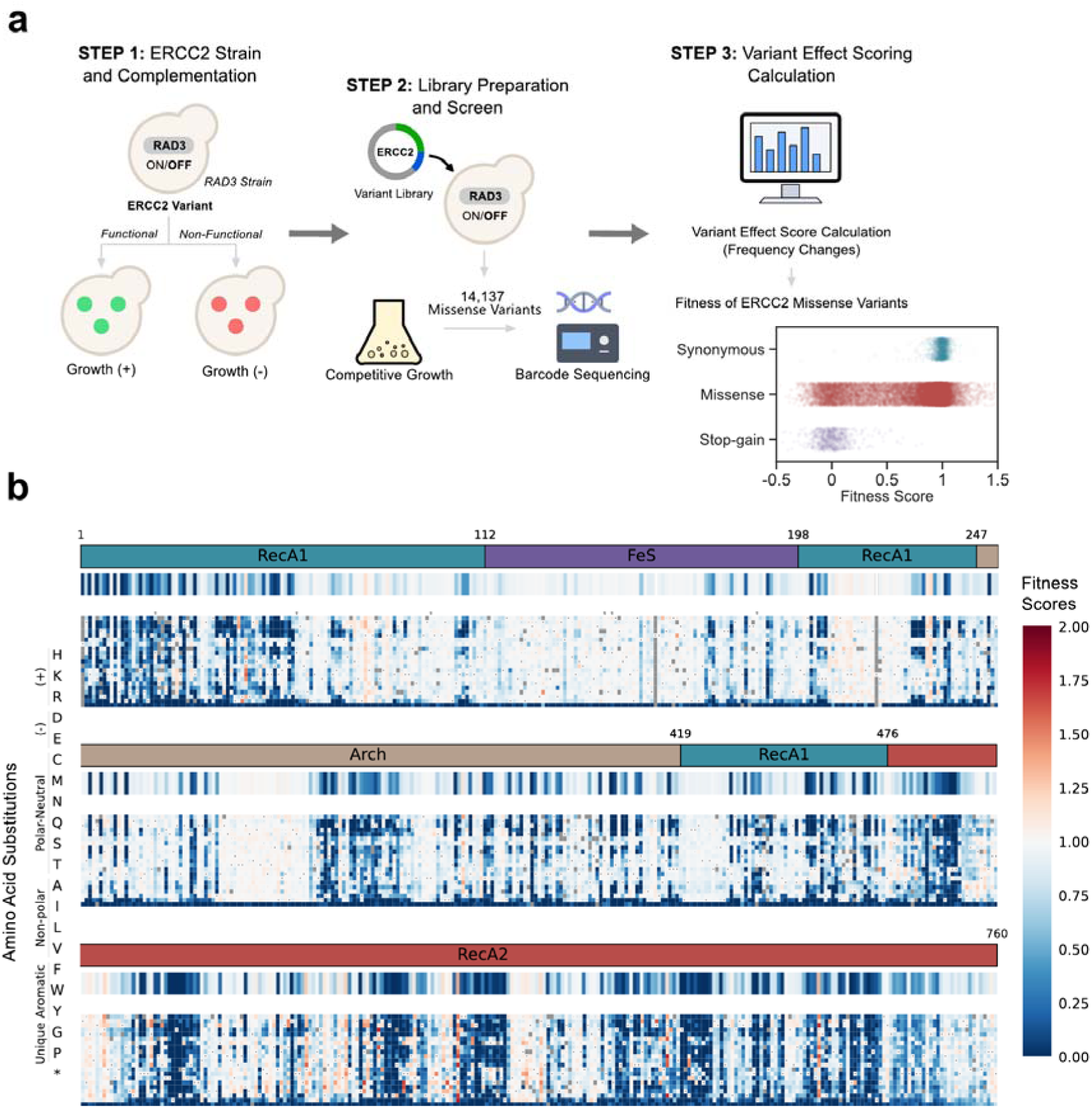
The fitness landscape of human *ERCC2* derived from yeast functional complementation. **(a)** Experimental workflow for deep mutational scanning of human *ERCC2*. **Step 1:** A TetO7-RAD3 yeast strain is used to assess complementation by human XPD; functional *ERCC2* variants restore growth, while non-functional variants impair it. **Step 2:** A saturation mutagenesis library of *ERCC2* is generated and introduced into yeast, followed by growth selection under defined conditions and barcode-based deep sequencing. **Step 3:** Sequencing data are processed to calculate variant effect scores from frequency changes, yielding quantitative fitness scores. Fitness scores for missense, synonymous, and stop-gain *ERCC2* variants are shown as the pipeline’s end point. **(b)** Heatmaps of human *ERCC2* variants aligned with domain annotations. The heatmap strip shows the median variant effect per position. The XPD amino acid sequence is divided from N- to C-terminus into three regions: upper (residues 1-253), middle (254-506), and lower (507-760). Colour coding indicates variant effect (fitness scores): dark blue, damaging; white, wild-type-like; red, hyper-complementing. Amino acid substitutions labelled on the left indicate the substitution order used across all maps.

Doxycycline treatment strongly reduced growth of the TetO7-*RAD3* strain, consistent with *RAD3* being essential, although residual slow growth indicated incomplete repression (Fig. S1b). Expression of wild-type *ERCC2* largely restored growth under *RAD3*-repressed conditions, consistent with previous evidence that human *ERCC2* can complement yeast *RAD3* (Fig. S1c). In the absence of doxycycline, XPD expression did not substantially alter growth, indicating that XPD has no additional beneficial or toxic effect when coexpressed with *RAD3*.

We next generated a plasmid-based saturation mutagenesis library covering approximately 98% of XPD single amino acid substitutions, represented by 350,813 unique barcodes (∼61% missense or stop-gain variants, 4% synonymous variants, and 33% wild-type; Fig. S2a). After growth selection under *RAD3*-repressed conditions, barcode enrichment was used to assign fitness scores to nearly all variants, scaled so that the median wild-type score was 1 and the median stop-gain score was 0. Biological replicates were highly concordant (Spearman ρ = 0.94; Fig. S2c), and synonymous and stop-gain variants formed the expected wild-type-like and loss-of-function distributions, respectively (synonymous mean = 0.99 ± 0.085; stop-gain mean = 0.00 ± 0.142). Missense variants showed a broad distribution of effects (mean = 0.69 ± 0.398), indicating that the screen captured a wide range of functional consequences. Damaging missense variants were distributed across XPD, but intolerance varied substantially between domains (Fig. 1b; Fig. S2d). The FeS domain was relatively tolerant to missense substitutions (mean fitness = 0.89), whereas RecA1, Arch and RecA2 showed lower mean fitness values of 0.70, 0.72 and 0.60, respectively.

### Growth-based complementation selectively reports transcription-associated XPD function

XPD has distinct roles in NER and transcription initiation. During NER, XPD helicase activity, DNA binding and ATPase-driven conformational changes are required for DNA repair.

During transcription initiation, however, XPD helicase activity is largely dispensable, and its essential role is thought to be structural, supporting TFIIH integrity and interactions with partner subunits^2,5,7^. Because NER is not required for yeast growth in the absence of exogenous DNA damage, we expected growth complementation to preferentially report transcription-associated rather than NER-specific XPD function.

To test this, we compared *ERCC2* fitness scores with previously reported transcription and DNA repair activities for 13 variants^2,5^. Fitness scores tracked transcriptional activity but not NER activity (Fig. 2a). Variants that impair DNA repair while retaining transcriptional function, including K48R, C134S, C155S, Y158A, F161A, F193A, R196A, R196E, K370A and K370E, showed near wild-type fitness. By contrast, variants that impair transcription-associated XPD function, including R722W and R324S, showed reduced fitness. The functionally neutral L372A variant, which retains function in both NER and transcription, yielded near-wild-type fitness. These results indicate that growth complementation primarily reports transcription-associated XPD function, rather than NER-specific activity.

**Figure 2:**
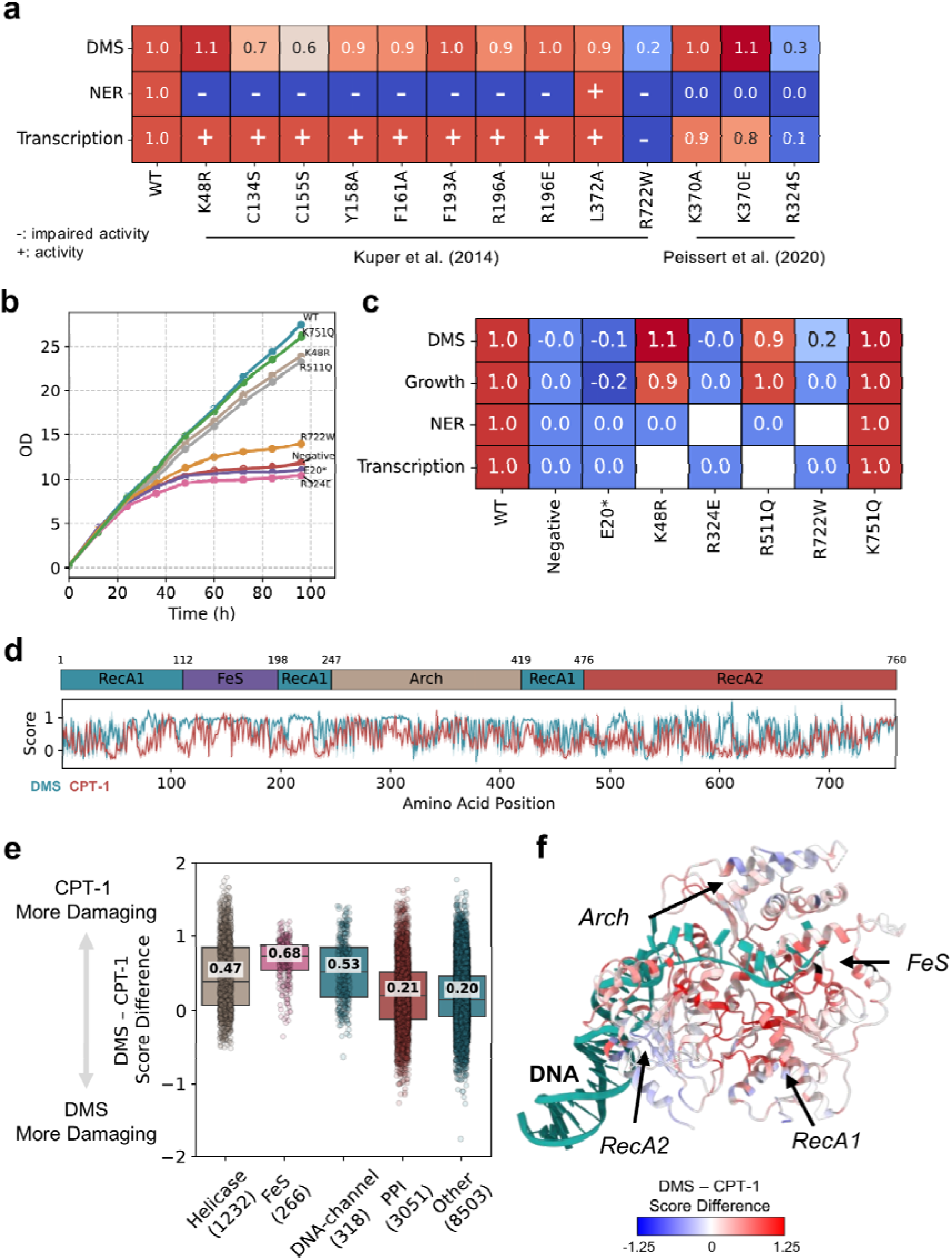
Mechanism selectivity of *ERCC2* fitness scores and their divergence from computational predictions. **(a)** Correspondence between growth fitness scores from DMS-based growth assays and previously reported NER and transcription initiation activities. Data from Kuper et al. and Peissert et al. are shown and labelled on the graph. Activity profiles from Kuper et al. are coded as - (no activity) or + (activity) as reported in the original publication, while Peissert et al. data are shown as numeric values. **(b)** OD600 measurements for eight TetO7-RAD3 strains: WT, K751Q, K48R, R511Q, R722W, Negative, E20*, and R324E. The WT strain expresses wild-type human XPD, while the Negative strain lacks XPD expression. **(c)** Correspondence between growth doubling times from panel (b) and DMS scores, consistent with literature-reported trends. **(d)** Comparison of fitness scores derived from DMS and CPT-1 predictions for *ERCC2* variants. Red indicates CPT-1 predictions and blue indicates DMS-derived fitness scores, each averaged per residue across all single amino acid substitutions; shaded regions show 95% confidence intervals (auto-calculated via Seaborn v0.13.2). Protein domains are shown above, aligned with amino acid position. Higher scores correspond to wild-type-like activity, while lower scores indicate more damaging effects. CPT-1 prediction scores were normalised to align with the DMS data (see Methods) **(e)** Differences between DMS and CPT-1 scores for functionally important residues in human XPD. Residues are grouped into five categories: Helicase, comprising conserved helicase motifs I, Ia, II, III, IV, V, and VI; FeS residues forming the FeS cluster pocket; DNA-channel, comprising residues contacting DNA in the 7AD8 PDB structure; PPI, comprising residues interacting with MAT1, XPB, p44, or p62 in the 6NMI PDB structure; and Other, comprising all other residues. Residues assigned to more than one category (e.g., overlapping PPI and Helicase residues) were excluded from the plot. Normalised scores (as in panel d) for DMS and CPT-1 were subtracted; mean values for each category are indicated (see Methods). **(f)** Structural mapping of DMS-CPT-1 differences onto the DNA-bound XPD structure (PDB: 7AD8), highlighting domains and the DNA channel. Blue indicates residues where the experimental DMS data show a more damaging effect than CPT-1 predictions; red indicates the opposite.

We confirmed this using individual growth assays for selected variants. The loss-of-function control E20* showed strongly impaired growth, while the benign variant K751Q grew similarly to wild-type (Fig. 2b, c). The transcription-defective variants R722W and R324E showed severe growth defects, whereas the DNA repair-defective variants K48R and R511Q showed near wild-type growth, consistent with their reported effects on DNA repair rather than transcription^1,2^.

We next compared DMS fitness scores with CPT-1^18^, a missense VEP based on sequence- and structure-derived information that has shown top performance in recent studies^19,20^. As in our recent comparison of discordance between MAVEs and VEPs^21^, we reasoned that divergences between experimental and computational variant effect scores could reveal classes of variants affecting molecular functions that are differentially captured by the two approaches. Unlike the growth assay, CPT-1 is not tied to a specific cellular readout and is therefore expected to reflect deleterious effects across a broader range of XPD functions. At the domain level, the largest discrepancies occurred in RecA1 and FeS, with additional differences in Arch and RecA2 (Fig. 2d). Analysis of specific functional features showed the greatest disagreement for FeS residues, followed by DNA-channel residues and conserved helicase motifs, whereas protein-protein interaction residues showed substantially stronger agreement (Fig. 2e). Mapping these discrepancies onto DNA-bound XPD showed that variants near the DNA channel and ATPase site were often predicted to be damaging by CPT-1 but retained near-wild-type growth in the DMS assay (Fig. 2f). These systematic patterns of agreement and disagreement distinguish molecular features that are strongly represented in the growth-based readout from those affecting XPD functions that are less important for complementation, illustrating how DMS–VEP discordance can provide information about variant mechanism.

### Transcription-associated fitness defects localise to XPD scaffold interfaces

Within TFIIH, XPD interacts with several partner subunits, including MAT1, p44, XPB and p62 (Fig. 3a). These interactions have distinct functional roles. p44 anchors XPD within the core TFIIH complex, while MAT1 links core TFIIH to the CDK-activating kinase (CAK) complex, enabling CDK7-mediated phosphorylation during transcription initiation. XPB and p62 also participate in interactions associated with DNA opening and regulation of XPD helicase activity during NER^22–24^.

**Figure 3:**
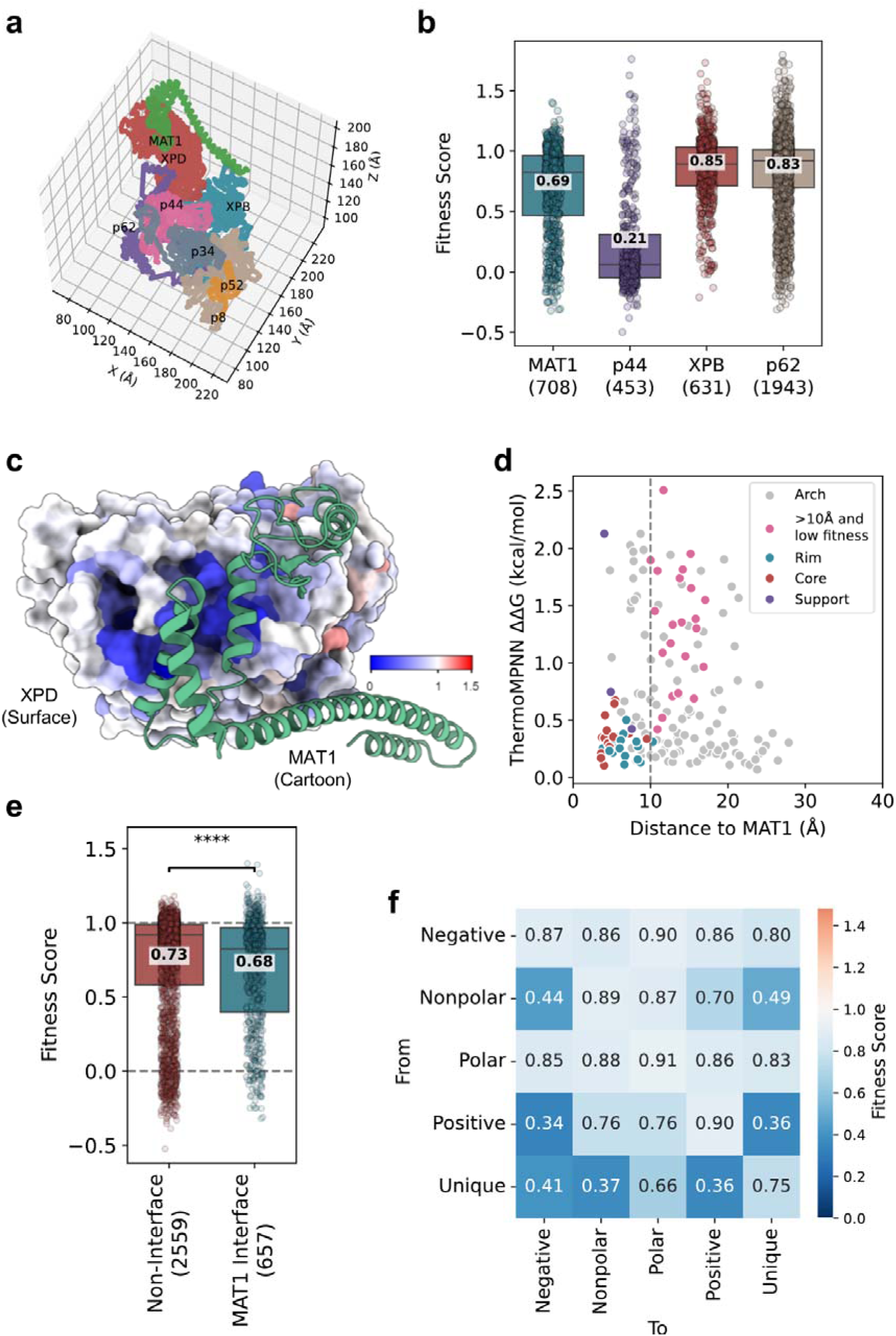
Interface-specific fitness effects and the structural basis of intolerance at the XPD-MAT1 interface.**(a)** 3D coordinate plot of the human TFIIH complex, showing constituent proteins MAT1, XPD, XPB, p44, p62, p34, p52, and p8. XPD’s interacting partners (MAT1, p44, XPB, and p62) are highlighted in distinct colours. **(b)** Distribution of growth fitness scores for residues at the protein-protein interaction interfaces with MAT1, p44, XPB, and p62. **(c)** Fitness scores mapped onto the 3D structure of XPD (PDB ID: 6NMI). XPD is shown as a surface coloured by mean fitness per residue, while MAT1 is depicted as a green cartoon. The colour bar indicates dark blue = damaging, white = wild-type-like, and red = hyper-complementing variants. **(d)** Relationship between ThermoMPNN ΔΔG (kcal/mol) and distance to MAT1 for Arch residues. MAT1 interface residues are coloured red (rim), teal (support), and purple (core). Residues >10 Å away from MAT1 that nevertheless show lower fitness (<0.5) are highlighted in pink; other residues in the Arch domain are shown in grey. **(e)** Distribution of fitness scores for Arch domain residues, comparing non-interface residues to those interacting with MAT1. Non-interface residues show significantly higher fitness compared with MAT1-interacting residues (Mann-Whitney U test, *p* = 2.279e-06). **(f)** Matrix of fitness scores based on amino acid substitution type (negative (D, E), non-polar (A, V, I, L, F, Y, W), polar (M, C, S, T, N, Q), positive (R, H, K), or unique (G, P)), with colour indicating resulting fitness scores.

Fitness scores differed markedly between interfaces (Fig. 3b). Variants at p44-contacting residues were particularly deleterious (mean fitness = 0.21), followed by those at the MAT1 interface (0.69), whereas variants at XPB- and p62-contacting residues had higher mean fitness scores of 0.85 and 0.83, respectively. This pattern is consistent with the growth assay predominantly reporting transcription-associated scaffold function, in which disruption of p44 anchoring or MAT1 recruitment can impair TFIIH integrity and transcriptional competence.

We next examined the structural basis of the fitness effects associated with the MAT1 interface. MAT1 binds primarily to the Arch domain of XPD, and mapping fitness scores onto the XPD-MAT1 structure revealed clusters of low-fitness variants within Arch-domain helices that directly contact MAT1, including R253-T276, A326-R345, P354-R364 and K370-R373 (Fig. 3c). However, strong intolerance was not restricted to the immediate interaction surface. Several regions more than 10 Å from MAT1, including C375-T385 and L391-V417, also contained low-fitness variants.

We therefore asked whether these distal effects might arise through destabilisation of XPD rather than direct disruption of MAT1 binding. We calculated the predicted change in protein folding free energy upon mutation (ΔΔG), where larger positive values indicate greater predicted destabilisation. Many low-fitness substitutions distal to MAT1 were associated with elevated predicted ΔΔG values (Fig. 3d), consistent with some of these effects being mediated by destabilisation of the Arch-domain structure. Consistent with the structural mapping, direct comparison of MAT1-contacting residues with other residues in the Arch domain showed a modest but significant reduction in fitness at the interface (mean fitness = 0.68 versus 0.73; Fig. 3e). Together, these results suggest that reduced fitness around the MAT1-binding region can arise through both direct disruption of the XPD-MAT1 interface and indirect destabilisation of the structural environment supporting it.

Patterns of amino acid substitution further suggested that local physicochemical compatibility contributes to MAT1-associated function. Changes involving proline and glycine were frequently poorly tolerated, consistent with structural constraints, while substitutions of positively charged residues were particularly deleterious when they introduced negative charge (Fig. 3f). This pattern is consistent with the importance of the positively charged XPD surface at the MAT1 interface.

Given the particularly strong intolerance of the p44 interface, we next investigated its structural basis in greater detail. p44 contacts XPD primarily through the RecA2 domain, and mapping fitness scores onto the XPD-p44 structure revealed pronounced intolerance across the interaction surface (Fig. 4a). Direct comparison with other RecA2 residues confirmed that p44-contacting residues were substantially less tolerant to substitution (mean fitness = 0.21 versus 0.64; Fig. 4b), consistent with the central role of p44 in anchoring XPD within core TFIIH.

**Figure 4:**
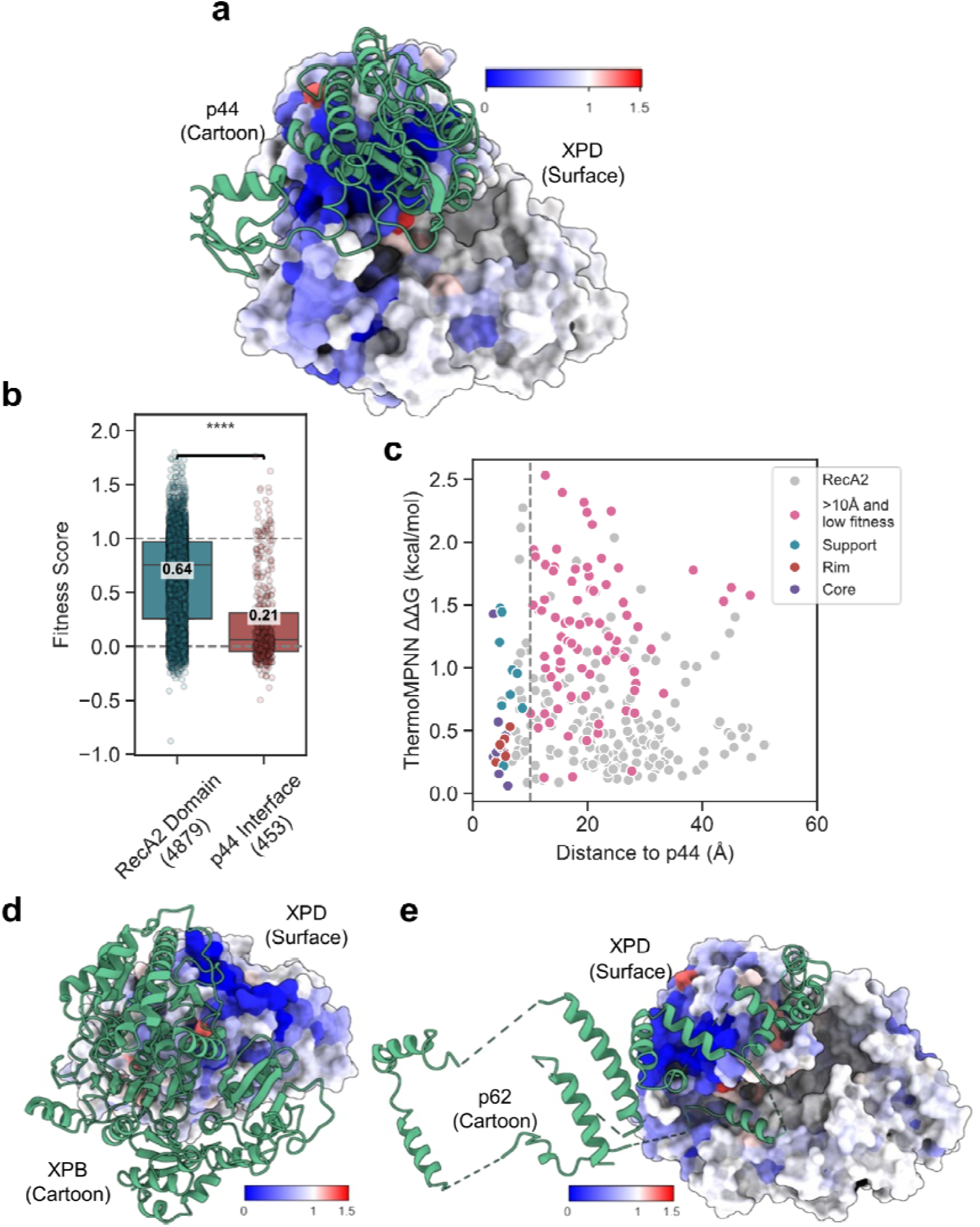
Structural determinants of fitness effects at the XPD-p44, XPD-XPB and XPD-p62 interfaces. **(a)** Fitness scores mapped onto the 3D structure of XPD (PDB ID: 6NMI). XPD is shown as a surface coloured by mean fitness per position, with p44 depicted as a green cartoon. **(b)** Distribution of fitness scores for RecA2 domain residues, comparing non-interface residues to those interacting with p44. Non-interface residues show significantly higher fitness compared with p44-interacting residues (Mann-Whitney U test, *p* = 7.481e-82). **(c)** Relationship between ThermoMPNN ΔΔG (kcal/mol) for RecA2 residues and their distance to p44. p44 interface residues are coloured red (rim), teal (support), and purple (core). Residues >10 Å away from p44 that nevertheless show lower fitness (<0.5) are highlighted in pink; other residues in the RecA2 domain are shown in grey. **(d-e)** Fitness scores mapped onto XPD 3D structures interacting with XPB (d) and p62 (e). Colour mapping is consistent with the p44 and MAT1 visualisations.

As observed for the MAT1-binding region, low-fitness variants were not restricted to residues directly contacting p44. Several distal regions within RecA2, including β-sheet regions spanning L490-V599, showed pronounced intolerance despite lying more than 10 Å from the interaction surface. Many of these distal low-fitness substitutions also had elevated predicted ΔΔG values (Fig. 4c), consistent with destabilisation of the RecA2 structural core. Thus, reduced fitness around the p44-binding region appears to arise through both direct disruption of the XPD-p44 interface and indirect effects on the structural stability of RecA2.

Finally, structural mapping of the comparatively tolerant XPB and p62 interfaces showed little evidence of localised intolerance around either interaction surface (Fig. 4d, e). These results suggest that the XPD-XPB and XPD-p62 contact surfaces are less critical for the functions required for complementation under standard growth conditions. Together with the pronounced intolerance of the p44 interface and more modest effects at the MAT1 interface, this pattern further supports the conclusion that the assay preferentially captures transcription-associated scaffold functions of XPD rather than the full range of molecular activities required during NER.

### Mechanism-selective DMS distinguishes ERCC2 disease phenotypes

The distinct molecular functions of XPD are reflected in its associated disease phenotypes, with XP primarily linked to defects in NER and TTD more closely associated with disruption of transcription-associated TFIIH function. Given the mechanism selectivity of the complementation assay, we therefore asked whether DMS fitness scores also captured differences between variants associated with these two phenotypes.

We first considered the distribution of previously identified pathogenic *ERCC2* missense variants across XPD. These comprised 18 variants associated with XP, 16 associated with TTD, and 9 variants that have been associated with both disease phenotypes (XP/TTD). All three classes were distributed across much of XPD and showed a common enrichment in the C-terminal RecA2 domain (Fig. 5a), indicating that disease phenotype cannot be explained simply by broad domain localisation. However, several phenotype-associated regional differences were also apparent. Variants within the three discontinuous RecA1 segments were observed only among XP and XP/TTD cases, while TTD-associated variants were absent from these regions. Conversely, variants within the Arch domain were observed only among TTD cases. These patterns are consistent with the distinct molecular functions underlying the two disease phenotypes: RecA1 contributes directly to the helicase and NER activities of XPD, whereas the Arch domain has an important structural role in transcription-associated TFIIH function, including interaction with MAT1. Thus, although XP- and TTD-associated variants overlap substantially in their overall structural distribution, their regional localisation shows differences that are consistent with the separable NER and transcription-associated functions of XPD.

**Figure 5:**
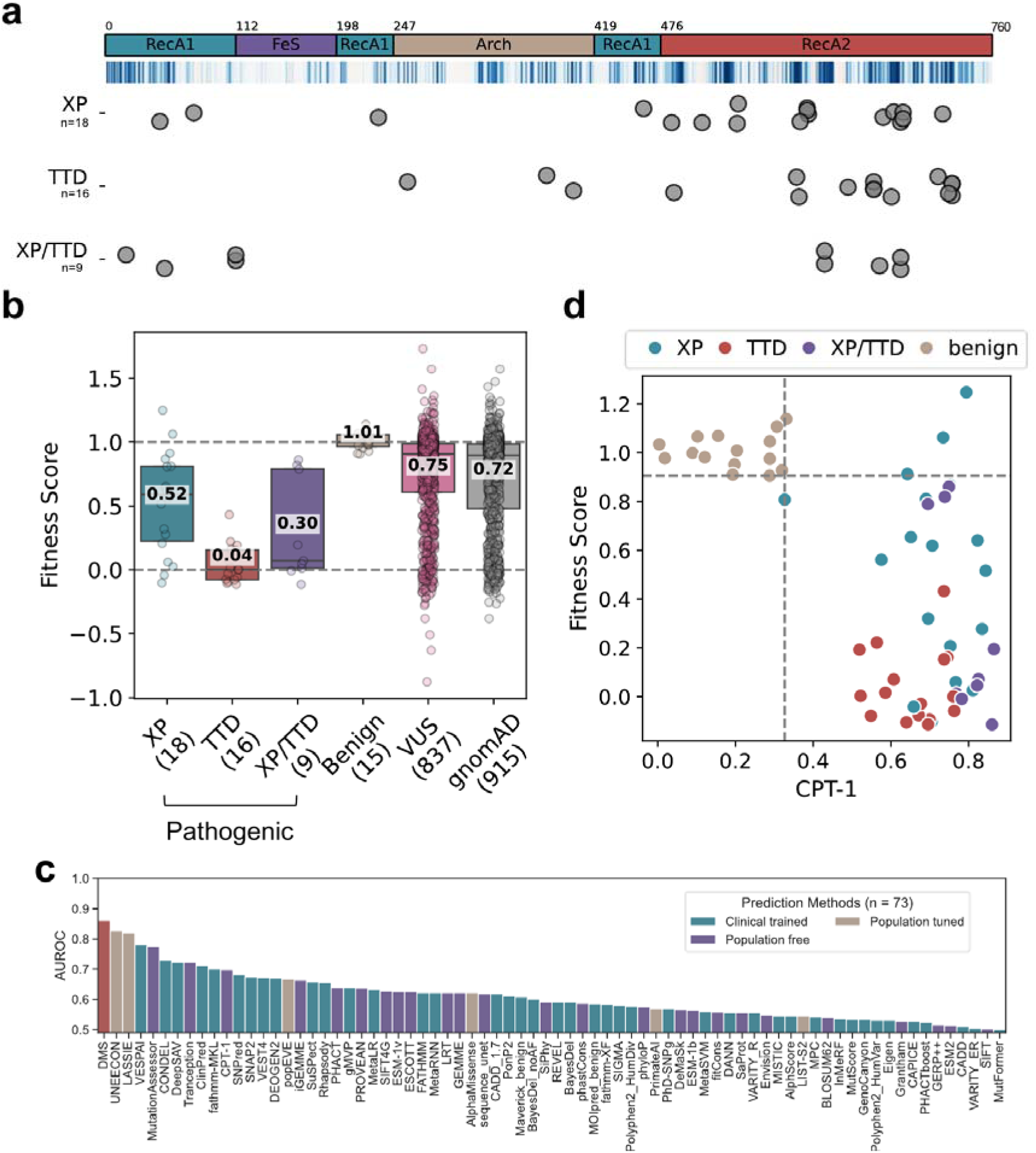
DMS fitness scores distinguish disease phenotypes associated with *ERCC2*. **(a)** XPD domain annotations with per-position disease-causing variants. Dots below domain annotations indicate disease-causing variants in three phenotype categories: XP (18), TTD (16), and XP/TTD (9). Data were collected from the literature (see Methods for details). **(b)** Fitness score distributions for *ERCC2* variants across six groups: XP, TTD, XP/TTD, benign, VUS (ClinVar and LOVD), and gnomAD. Variant counts for each subgroup are indicated in parentheses. Of the 848 VUS and 930 gnomAD variants, 837 and 915, respectively, were visualised, as only these variants had DMS data available. **(c)** AUROC values for discrimination between TTD- and XP-associated ERCC2 variants across 73 VEPs and DMS. **(d)** Scatter plot of *ERCC2* disease-causing variants, with DMS scores on the y-axis and CPT-1 scores on the x-axis. Variants are coloured by category: XP, TTD, XP/TTD, and benign.

Given the transcription-associated selectivity of the complementation assay, we expected TTD-associated variants to show lower fitness than XP-associated variants, some of which primarily impair NER. Consistent with this, TTD variants had markedly reduced fitness scores (mean = 0.04) compared with XP variants (mean = 0.52; Fig. 5b). Variants associated with both XP and TTD phenotypes (XP/TTD) showed intermediate fitness scores (mean = 0.30), although this group was smaller. In contrast, known benign variants showed wild-type-like fitness (mean = 1.01), while VUS from ClinVar and LOVD and population variants from gnomAD had lower mean fitness scores of 0.75 and 0.72, respectively, with broad distributions.

We next assessed the ability of our DMS fitness scores to distinguish between pathogenic and benign missense variants, as measured by the area under the receiver operating characteristic curve (AUROC). Consistent with the score distributions shown in Fig. 5b, fitness scores showed complete separation (AUROC = 1) between TTD and benign variants, and between XP/TTD and benign variants, with all TTD and XP/TTD variants having lower fitness values than all of the benign variants. Discrimination was lower for XP, reflecting the greater overlap between pathogenic and benign variants, but remained high (AUROC = 0.90). Importantly, this reduced discrimination for XP was asymmetric: it reflected a subset of XP-associated variants with near wild-type fitness rather than benign variants with low fitness. Low fitness therefore remained highly informative for XP, whereas high fitness did not exclude pathogenicity.

We also compared our DMS fitness scores with 73 missense VEPs (Fig. S3). In contrast to the DMS, VEPs generally showed similarly strong pathogenic-versus-benign discrimination across TTD, XP and XP/TTD variants, with multiple predictors achieving perfect discrimination in the individual comparisons. VEP performance was therefore much less dependent on whether pathogenicity arose through transcription-associated or NER-associated XPD dysfunction. However, two caveats apply to these comparisons. First, the reference sets are small, including only 15 benign variants, so many methods approach the upper limit of performance and differences among the strongest predictors should not be over-interpreted. Second, many of the VEPs are trained using clinically labelled variants and may therefore have encountered some *ERCC2* variants, or closely related clinical information, during model development. Consistent with this possibility, clinically trained predictors comprise a substantial fraction of the highest-performing methods in Fig. S3, although several population-free predictors also perform strongly. Nonetheless, the overall contrast with the DMS is clear: computational predictors generally distinguished pathogenic variants from benign variation similarly across disease phenotypes, whereas DMS pathogenicity discrimination showed a pronounced phenotype dependence, with substantially stronger separation for TTD than for XP.

Although VEPs therefore performed strongly at distinguishing pathogenic from benign variation irrespective of phenotype, we next asked whether the scores could distinguish between the two major disease phenotypes themselves. Restricting the analysis to variants associated specifically with XP or TTD, DMS fitness scores discriminated between the two phenotypic classes with an AUROC of 0.86 (Fig. 5c), higher than all 73 VEPs tested. Several VEPs also showed substantial phenotype discrimination, but none matched the DMS.

The complementarity between DMS and computational predictors is illustrated by direct comparison of our fitness scores with CPT-1 (Fig. 5d). CPT-1 largely separates benign from disease-associated variants along the computational score axis, but shows little corresponding structure among the disease phenotypes, with TTD-associated variants tending towards slightly less damaging CPT-1 scores. In contrast, the DMS axis provides the phenotype-dependent information described above, with TTD-associated variants showing uniformly low fitness and XP variants showing much greater heterogeneity. Consequently, variants that are less clearly resolved by one score tend to be distinguished by the other: benign variants occupy a distinct region of the joint score space, while pathogenic variants are further stratified according to phenotype by DMS fitness. Thus, the joint score space provides complete separation of benign from disease-associated variants in this dataset, while also strongly distinguishing the major disease phenotypes, illustrating the complementary information provided by general variant effect prediction and a mechanism-selective MAVE.

### Clinical calibration of ERCC2 DMS scores using ACMG/AMP evidence strengths

Finally, we investigated whether DMS-derived fitness scores could be translated into clinically interpretable evidence. Fitness scores were calibrated using *acmgscaler*^25^, which estimates score-specific likelihood ratios from the distributions of known pathogenic and benign variants and maps these onto ACMG/AMP evidence strengths within a Bayesian framework^26^. The method uses bootstrapped kernel density estimation, adaptive regularisation and confidence bounds to provide robust evidence assignments across the score range. Previous benchmarking of *acmgscaler* showed that calibrations using as few as 10 pathogenic and 10 benign variants are generally conservative rather than prone to inflated evidence strengths^25^.

We calibrated DMS evidence separately for TTD (Fig. 6a) and XP (Fig. 6b), using the same benign reference variants but phenotype-specific sets of pathogenic missense variants.

**Figure 6:**
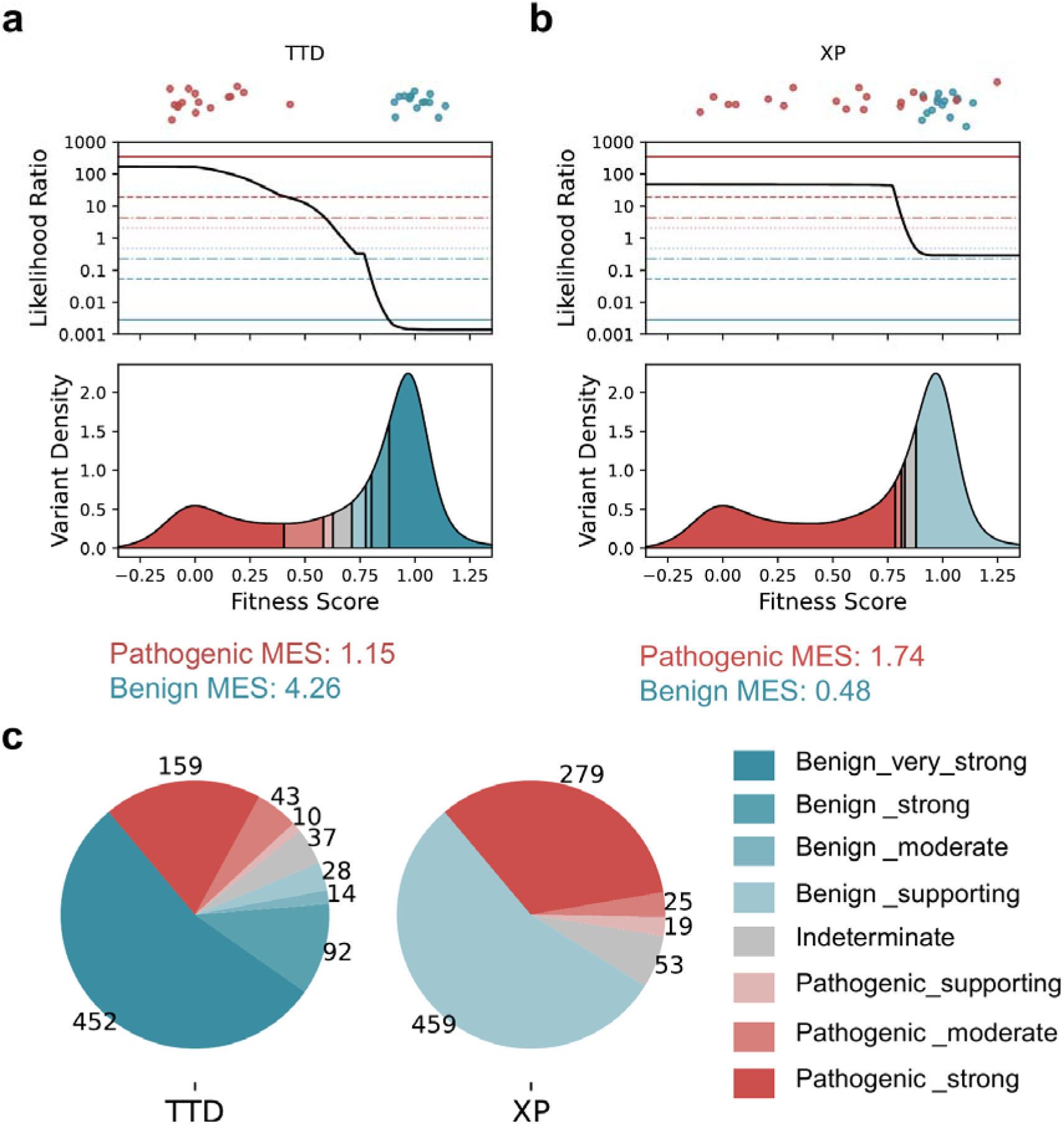
Phenotype-specific ACMG/AMP calibration of DMS fitness scores and its application to variants of uncertain significance. **(a-b) Top panel:** fitness score distributions for the reference datasets used to calculate likelihood ratios, with blue indicating the benign reference dataset and red the pathogenic reference dataset. **Middle panel**: likelihood ratios calculated using acmgscaler, with the x-axis showing DMS-derived fitness scores and the y-axis showing likelihood ratios. **Bottom panel:** distribution of ACMG/AMP evidence strengths across all measured variants; horizontal lines indicate evidence strength thresholds. Panel (a) corresponds to the TTD phenotype; panel (b) corresponds to the XP phenotype. **(c)** Total ACMG evidence criteria assigned to ClinVar and LOVD VUS, separated by TTD and XP phenotype. The number of variants in each category is indicated. 835 of 848 VUS were visualised, as only these had DMS data from both replicates.

Consistent with the much stronger separation between TTD and benign fitness scores, the resulting calibrations differed markedly between the two phenotypes, particularly for benign evidence. For TTD, all benign variants were tightly clustered at high fitness scores and clearly separated from pathogenic variants, allowing the calibration to reach very strong evidence for benignity. In contrast, several XP-associated pathogenic variants retained high fitness scores overlapping the benign range, limiting benign evidence to supporting strength. This difference was reflected in mean evidence strength (MES), which summarises the average ACMG/AMP evidence points generated across a variant set^20^, with benign MES nearly nine-fold higher for TTD than for XP (4.26 versus 0.48).

For pathogenic evidence, both calibrations reached strong evidence strength because no benign variants occurred at low fitness scores. However, the pathogenic score distributions differed substantially. TTD-associated variants were tightly concentrated at very low fitness scores, producing much higher likelihood ratios than XP in this region, but relatively little pathogenic evidence at intermediate scores. XP-associated variants were more broadly distributed, so pathogenic evidence extended across a wider range of fitness scores. As a consequence, pathogenic MES was higher for XP than for TTD (1.74 versus 1.15). This does not indicate stronger pathogenic-versus-benign discrimination for XP, but instead reflects the broader range of fitness scores over which XP-associated variants provide pathogenic evidence.

To illustrate the utility of this approach, we applied the calibrated scores to 835 missense variants classified as VUS in ClinVar and LOVD (Fig. 6c; Table S2). Using the TTD calibration, 95.6% of VUS received at least supporting evidence of pathogenicity or benignity, including 54.1% with very strong benign evidence and 19.0% with strong pathogenic evidence. For XP, 54.9% received supporting benign evidence and 33.4% received strong pathogenic evidence. The clinical utility of the assay therefore differs substantially between the two phenotypes. For TTD, DMS scores provide useful evidence in both directions, helping both to identify potentially pathogenic variants and to exclude variants unlikely to contribute to disease. For XP, the assay is primarily informative for pathogenic evidence: low fitness can provide strong support for pathogenicity and potentially contribute to diagnosis, whereas high fitness provides only limited evidence against pathogenicity because NER-specific defects may retain near wild-type growth. Thus, mechanism-aware calibration allows the same functional dataset to provide clinically useful evidence while accounting for the disease mechanism relevant to each phenotype.

## Discussion

Our results show that yeast complementation DMS provides a mechanism-selective map of *ERCC2* variant effects. Rather than measuring all XPD functions equally, the growth-based assay primarily reports the subset of XPD activities required to complement yeast *RAD3* under standard growth conditions. Variants affecting helicase activity, ATPase function, DNA binding and NER-associated regions such as RecA1 and FeS often retained near wild-type fitness, whereas p44-contacting residues showed pronounced intolerance and more modest effects were observed at the MAT1 interface. These interactions are central to TFIIH stability and transcriptional function: p44 anchors XPD within TFIIH, while MAT1 connects the core complex to the CDK-activating kinase subcomplex^5,7,27^. Importantly, low-fitness variants around these interaction surfaces were not restricted to residues directly contacting p44 or MAT1. Distal substitutions with low fitness were frequently predicted to destabilise XPD, suggesting that disruption of transcription-associated scaffold function can arise through both direct loss of intermolecular contacts and indirect destabilisation of the structural environment supporting them. Thus, the assay defines at variant resolution which regions and classes of *ERCC2* variation contribute to the transcription-associated functions required for complementation, rather than providing a universal measure of XPD dysfunction.

This mechanism selectivity is a limitation only when fitness scores are treated as universal measures of pathogenicity; interpreted in the appropriate biological context, it instead provides mechanistic information. This was particularly apparent when we compared DMS scores with computational VEPs. VEPs generally showed excellent discrimination between pathogenic and benign *ERCC2* variants irrespective of disease phenotype, whereas DMS fitness scores distinguished XP- from TTD-associated variants better than any of the 73 VEPs tested. Systematic disagreement between experimental and computational scores therefore need not simply represent measurement error, but can reflect differences in the biological information captured by the two approaches. Computational VEPs provide broad information about whether a substitution is likely to disrupt protein function, whereas the DMS readout additionally reports whether that disruption affects the transcription-associated functions required in the complementation assay. More generally, functional assays of multifunctional proteins should therefore be interpreted in terms of the specific molecular activities required by the assay, rather than judged solely by their agreement with broad measures of variant deleteriousness. Together, experimental and computational approaches can provide complementary information about pathogenicity and disease mechanism that neither captures as effectively alone.

The phenotype-associated fitness distributions also emphasise that XP and TTD do not represent completely discrete molecular categories. TTD-associated variants were strongly concentrated at low fitness scores, XP variants showed a much broader distribution, and variants reported with both XP and TTD phenotypes tended to show intermediate effects. Some XP-associated variants therefore perturb functions shared with transcription-associated disease mechanisms, whereas others appear to produce more selective NER defects that are less apparent under standard growth conditions. This complexity is consistent with previous reports of overlapping XP and TTD phenotypes^28,29^, and is also expected in a recessive disease context, where clinical outcome may depend on the combination of alleles, their residual activities, genetic background and ascertainment of specific clinical features. Rather than discrete categories, the phenotype-associated fitness distributions suggest a graded relationship between variant effect and disease mechanism, reflecting differences in the extent to which individual variants disrupt the transcription-associated functions captured by the assay.

Calibration of the DMS scores within an ACMG/AMP framework showed how this mechanism selectivity can be incorporated into clinical interpretation. For TTD, the strong separation between pathogenic and benign variants meant that DMS scores provided substantial evidence in both directions, supporting pathogenicity for low-fitness variants while also providing strong evidence against pathogenicity for variants with preserved fitness. However, because few TTD pathogenic or benign variants fall at intermediate fitness scores, evidence assigned in this range should be interpreted cautiously. For XP, the clinical utility was more asymmetric. Low fitness remained strongly associated with pathogenic variation and could therefore contribute useful evidence towards diagnosis, but preserved fitness provided much weaker evidence against pathogenicity because variants with selective NER defects can retain near-wild-type growth. This illustrates why phenotype-specific calibration is more appropriate than applying a single pathogenicity threshold across all *ERCC2*-associated disease. In practice, these phenotype-specific calibrations would be applied in the context of a clinical suspicion of XP or TTD, rather than used independently to infer the disease phenotype.

The TTD calibration assigned ‘very strong’ benign evidence to a substantial fraction of variants. Within the Bayesian ACMG/AMP framework, evidence of this magnitude would by itself be sufficient to cross the threshold for a benign classification, making these assignments potentially clinically consequential. However, in our previous validation of *acmgscaler*, we specifically noted that the ‘very strong’ evidence category warrants additional caution, as its performance is more difficult to assess because of data sparsity and occasional instability in likelihood-ratio estimates^25^. We therefore consider it prudent, for current clinical application, to cap *acmgscaler*-derived evidence at strong until very strong evidence assignments can be validated more extensively using larger and independent reference datasets.

Several limitations remain. Yeast complementation may not capture human-specific regulatory effects, post-translational modifications or all TFIIH interactions, and standard growth conditions do not directly challenge NER activity. Incomplete repression of endogenous *RAD3* may also partially buffer hypomorphic *ERCC2* variants, causing some intermediate-effect variants to appear closer to wild type. Future assays incorporating DNA-damaging conditions, tighter *RAD3* depletion or complementary mammalian systems could therefore extend the functional map to variants that selectively affect DNA repair. Supporting the feasibility of this approach, a recent study profiled a panel of cancer-associated *ERCC2* variants using CRISPR-Select in human MCF10A cells^30^; as expected, variants that disrupt the NER pathway were found to sensitise cells to cisplatin treatment. Clinical calibration is additionally constrained by the relatively small number of confidently classified *ERCC2* variants and by uncertainty in assigning individual alleles to specific disease phenotypes, particularly in compound heterozygous patients. Evidence strengths should therefore be interpreted alongside other clinical and molecular data. Despite these limitations, this XPD map shows how a mechanism-selective MAVE can connect variant effects to distinct disease mechanisms and support phenotype-aware clinical interpretation.

## Materials and Methods

### Preparation of the Saccharomyces cerevisiae TetO7-RAD3 Strain

We generated a yeast strain that enables controlled expression of *RAD3*, the yeast orthologue of human *ERCC2*. *RAD3* expression was regulated using a Tet-off system, in which the addition of doxycycline (10-20 µg/mL) effectively represses transcription from Tet-responsive promoters^31,32^. To achieve this, we replaced the native *RAD3* promoter by homologous recombination using a custom-designed integration cassette (Fig. S1a). The cassette consisted of a synthetic TetO7 promoter, a geneticin (G418 sulfate) selectable marker, and homologous arms flanking the target site: a left arm spanning positions −324 to - 264 and a right arm spanning positions −5 to +55 relative to the start codon. The cassette was integrated into the Yeast Tet-promoters Hughes parental strain R1158 (URA3::CMV-tTA MATa his3-1 leu2-0 met15-0) using the standard LiAc/SS carrier DNA/PEG method^33^. This strain contains the Tet transactivator gene integrated into its genome, thereby enabling control of the TetO7 promoter through doxycycline treatment. Positive transformants were selected, and correct integration and cassette sequence at the target locus were verified.

### Functional Complementation of Yeast RAD3 by Human ERCC2 in the TetO7-RAD3 Strain

Human *ERCC2* cDNA (UniProtKB: P18074) was codon-optimised for *Saccharomyces cerevisiae* using the IDT Codon Optimization Tool and expressed from the pMP1_XL plasmid in the TetO7-*RAD3* background. The pMP1_XL–*ERCC2* construct and the empty pMP1 vector (negative control) were introduced by LiAc/SS carrier DNA/PEG transformation. Positive transformants were cultured for 48 h with medium refreshed every 12 h and supplemented with doxycycline to repress endogenous *RAD3* expression.

To assess functional complementation, optical density (OD600) of each culture (three replicates per condition) was measured every 30 minutes for approximately 36 h using a Tecan Infinite M200 Pro plate reader. Growth rates were calculated by fitting a logistic (sigmoidal) growth model^34^ to the OD600 data using the equation:

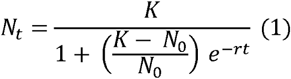

where is the OD600 at time , is the initial population size, is the carrying capacity, and is the intrinsic growth rate, reflecting the rate of cellular growth.

### Site-Directed Mutagenesis and Single-Variant Growth Analysis

We generated six specific *ERCC2* variants (E20*, K48R, R324E, R511Q, R722W, and K751Q) using the Q5® Site-Directed Mutagenesis Kit. E20* was designed as a complete loss-of-function variant; K48R and R511Q were associated with DNA repair defects while retaining transcription initiation activity; R324E and R722W were primarily linked to transcription defects; and K751Q was clinically reported as benign^1,2,5,35^.

Each variant was confirmed by whole-plasmid sequencing and transformed into the TetO7-*RAD3* strain. Single colonies of each variant were cultured in 5 mL medium, with ∼10-fold dilutions performed every 12 h across eight time points. Optical density at 600 nm (OD600) was measured at each interval, both before and after dilution. Doubling time was calculated as:

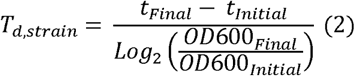

Where OD600_initisl_and OD600_final_ correspond to the starting and cumulative final optical densities (the latter corrected for serial dilution across the eight passages), andt_final_ and to the corresponding time points, respectively, such that _,initisl_ represents the time required for the population to double. Doubling times were normalised such that WT = 1 and the negative control = 0, consistent with the scaling used for DMS scores.

### Variant Library Preparation and Barcode Association

We generated a variant library using a plasmid-based saturation mutagenesis protocol^36^. This approach utilised customised oligonucleotide pool designs adapted from the SUNi mutagenesis protocol^37^. Systematic mutagenesis was performed at 65-amino-acid intervals, creating 13 defined segments to ensure uniform coverage. Each segment was cloned into the yeast expression plasmid pMP1_XL via Gateway LR recombination, with repeated reactions to enhance barcode diversity.

For sequencing, equal amounts of DNA from each segment were pooled and digested with HF-EcoRV and HF-SacI to excise the variant-containing region. The resulting fragments were purified and sequenced using the PacBio Revio system at Edinburgh Genomics.

Barcode-to-variant assignment was performed with the *alignparse* pipeline^38^, retaining high-quality, indel-free reads and generating consensus sequences from the filtered Circular Consensus Sequencing (CCS) data.

### Growth-based screen under normal conditions

We screened missense variants using a yeast-based growth assay. Preliminary experiments with wild-type and negative-control strains in the TetO7-*RAD3* background identified optimal conditions: 96 h total culture with ∼10-fold dilutions every 12 h. The screen was then performed under these conditions in two biological replicates, with samples collected at each time point and stored at −20°C.

Plasmids were extracted from each replicate and time point using the QIAprep Spin Miniprep Kit following pre-treatment with 1 M DTT and Zymolyase to enhance cell lysis. Barcodes were amplified using Phusion® High-Fidelity PCR with custom primers containing sequencing sites and unique 5-nt indices (16 cycles). PCR products (∼300 bp) were gel-purified, quality-checked on a BioAnalyzer, pooled in equimolar amounts, and sequenced on the Illumina NextSeq (P3) platform at the Wellcome Trust Sequencing Service.

### Data analysis and variant effect scoring

We used Enrich2 to calculate individual barcode scores from sequencing data, reflecting its support for barcode-based and time-series DMS experiments^39,40^. Barcode scores were then median-averaged to derive a single fitness score per variant. Based on prior optimisation, a 96-hour growth period with ∼10-fold dilutions every 12 h (8 time points) captured fitness differences between negative control and wild-type strains. Experimental noise across the time course (12–96 h) was evaluated as previously described^16^, showing that the full 96-hour duration minimised noise (Fig. S2b); hence, fitness scores from the 96-hour time point were used for further analysis.

Barcode scores were calculated with predetermined parameters (T0 ≥ 0, Max N: 0, MinQ: 20) for both replicates. For each replicate, fitness scores were aggregated either by median-averaging multiple barcodes or using the single barcode score if only one was present. Scores from biological replicates were combined into a single, robust variant score using a weighted mean via the DescrStatsW function from the statsmodels package. Weighted standard errors (WSE) were also calculated for each score.

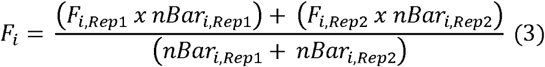

Where F_i_ is the weighted mean fitness score of variant ; Fi,_Rep_and Fi,_Rep_ are the fitness scores from biological replicates 1 and 2; and nBar_iRep1_and nBar_iRep_ are the corresponding number of barcodes.

### Structural and VEP Analysis

For structural analysis, we used two PDB structures: 6NMI and 7AD8. 7AD8 was employed solely to visualise the DNA-bound form, while 6NMI, which includes complete interfaces with MAT1, p44, and XPB and a partial interface with p62, was used to extract protein-protein interacting residues and for visualisation.

Solvent-accessible surface area (SASA) was calculated for residues using AREAIMOL. Residues with ≥25% surface exposure were defined as surface, and those with lower exposure as interior, using reference SASA values^41^. Interface residues were classified as core, rim or support, as previously defined^42^. Protein-protein interacting residues were extracted from PDB structure 6NMI. DNA-channel residues and FeS interacting residues were extracted from PDB structure 7AD8. Conserved helicase motifs were taken from a previous study^9^. Fitness scores were mapped onto PDB structures and visualised with ChimeraX^43^.

We also computed free energy changes (ΔΔG, kcal/mol) for missense variants using ThermoMPNN via its Google Colab interface, based on the 6NMI structure^44^. Scores for the 73 VEPs were obtained as part of our previous study^19^.

To facilitate comparison, DMS fitness scores and CPT-1 scores were normalised using clinically annotated benign variants (n = 15) and pathogenic variants associated with XP, TTD and XP/TTD (n = 43). For both datasets, scores were scaled such that the median benign variant score was 1 and the median pathogenic variant score was 0 (Fig. S2e). This normalisation using clinically annotated benign and pathogenic variants was used only for visual comparison between the two score sets and for identifying relative DMS vs CPT-1 discrepancies in Fig. 2d-f. All classification and calibration analyses used the original DMS and VEP scores, with no prior scaling to the clinical reference variants.

### Clinical Data Curation

We curated a dataset of 15 benign *ERCC2* missense variants from the ClinVar and LOVD databases^35,45^. For the pathogenic dataset, 43 missense variants were collected from the published literature^8,46–48^. Variant-specific phenotypes (XP, TTD, XP/TTD) were obtained from these source publications. We compiled 848 VUS from the LOVD and ClinVar databases, and 930 population variants from gnomAD v4.1. All data curation steps were completed as of October 2025. All variants, along with their associated disease phenotypes, are listed in Table S1.

### Area Under the Receiver Operating Characteristic Curve Analysis

We assessed the classification performance of our DMS data alongside VEPs, each providing scores for ≥95% of curated pathogenic and benign variants and included in a recent benchmark study^19^. Performance was evaluated using the AUROC, calculated with roc_auc_score() from sklearn.metrics via our custom roc_auc_calculate() function.

### Calculation of ACMG/AMP Evidence Strengths for TTD and XP subsets

We used *acmgscaler* to convert DMS-derived fitness scores into standardised ACMG/AMP evidence strengths for variant classification^25^, using the same variant sets as in the AUROC analysis. Datasets were formatted according to the tool’s requirements and processed via Google Colab, applying the default prior probability of 0.1 for calibration. To ensure robust evidence calculations, only *ERCC2* variants with fitness scores present in both replicate experiments were included; any variant detected in only a single replicate was excluded from the acmgscaler input.

## Acknowledgements

JAM was supported by funding from the European Research Council (ERC) under the European Union’s Horizon 2020 research and innovation programme (grant agreement No. 101001169) and by funding from the Medical Research Council (MRC) Human Genetics Unit core grant (MC_UU_00035/9).

## Data Availability

Fitness scores will be deposited at MaveDB and sequencing data at NCBI SRA prior to publication.

## Supplemental Figures

**Figure S1:**
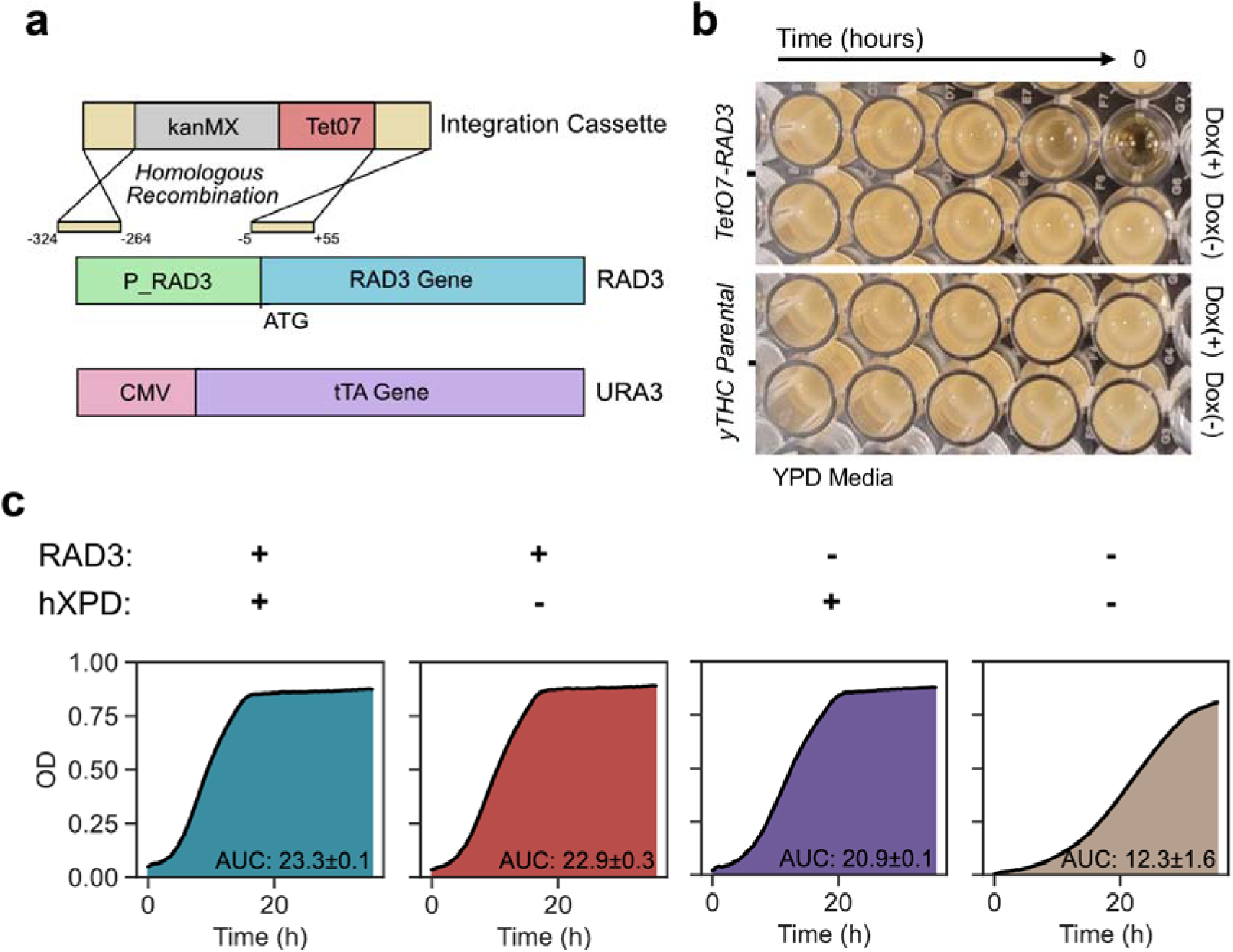
Generation, validation, and complementation of the TetO7-RAD3 strain. **(a)** Schematic of the integration cassette design and homologous recombination strategy used to generate the TetO7-RAD3 strain. **(b)** Functional validation of the TetO7-RAD3 strain, demonstrating effective repression of RAD3 expression in response to doxycycline treatment, with growth as the readout. **(c)** Functional complementation of yeast RAD3 by human *ERCC2*. The upper diagram depicts doxycycline-controlled expression of yeast RAD3 and human *ERCC2*, with expression states indicated (’+’, on; ’–’, off). Growth curves below correspond to the indicated expression conditions; growth rates were quantified as the area under the curve (AUC). Values represent the mean of three independent biological replicates.

**Figure S2:**
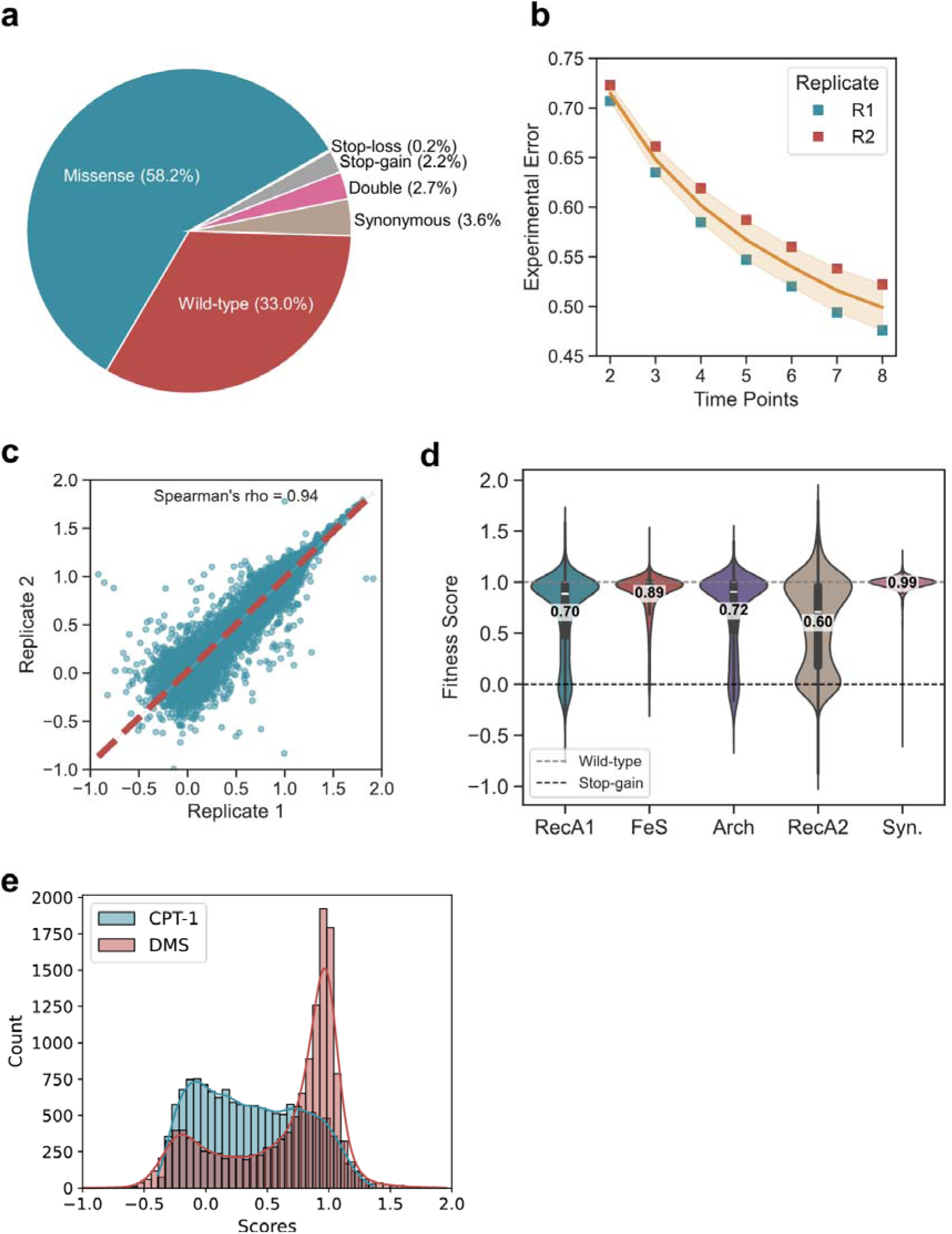
Variant library composition and fitness score quality.**(a)** Distribution of total barcodes assigned to each *ERCC2* variant category, shown as a pie chart: missense, wild-type, synonymous, double, stop-gain, and stop-loss. **(b)** Experimental error values across time points 2-8 for biological replicates. **(c)** Correlation of fitness scores between biological replicates 1 and 2. **(d)** Domain-specific distribution of fitness scores across XPD domains (RecA1, FeS, Arch, and RecA2). Synonymous (Syn.) *ERCC2* variants represent the wild-type-like phenotype. All domains differed significantly from synonymous variants (Mann-Whitney test, p < 0.05). **(e)** Distribution of normalised CPT-1 and DMS scores. Scores for *ERCC2* variants were normalised using clinically annotated benign (n = 15) and pathogenic variants associated with XP, TTD or XP/TTD (n = 43) reference sets (see Methods for details). Red indicates the DMS score distribution; blue indicates the CPT-1 score distribution after normalisation.

**Figure S3:**
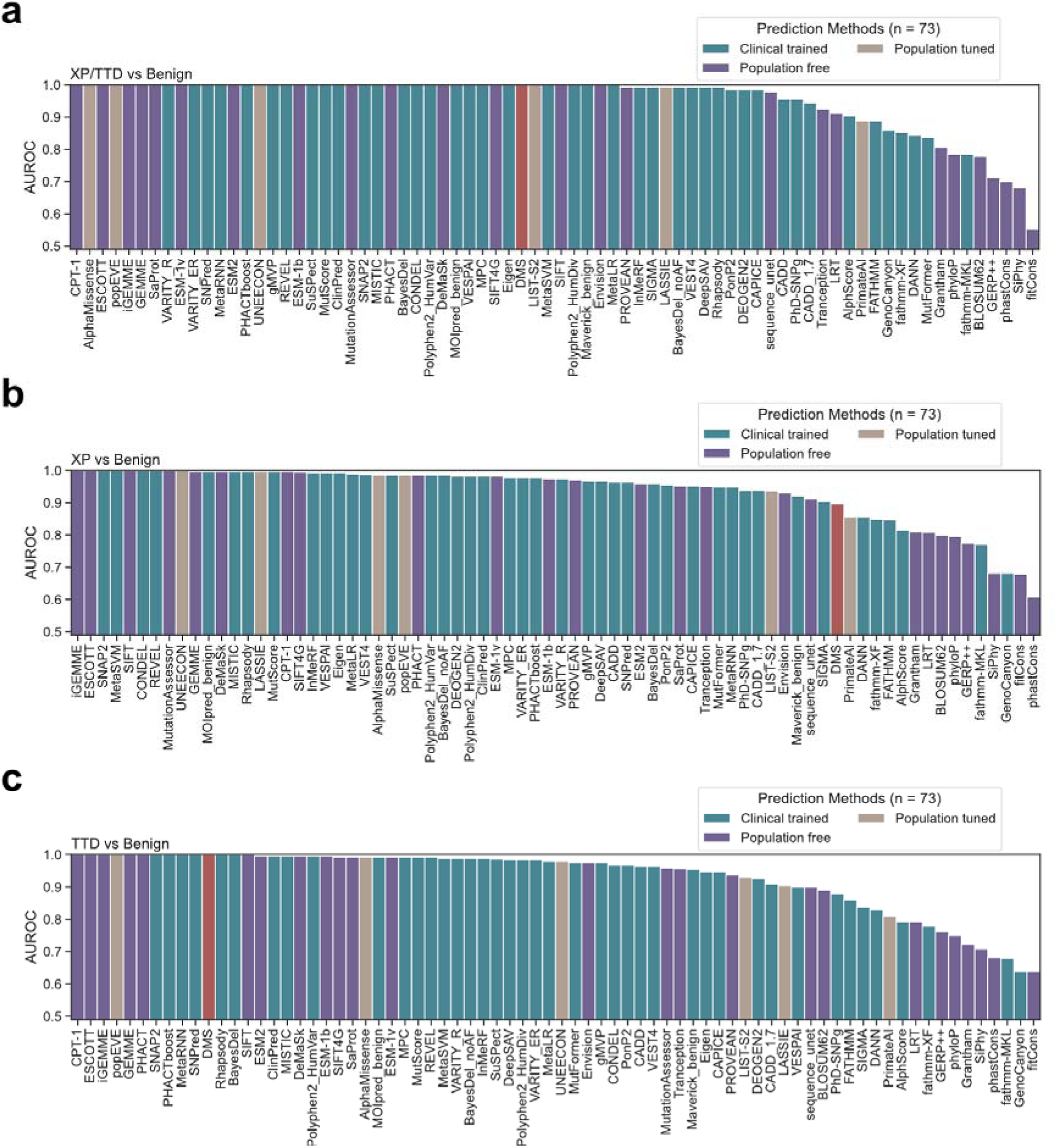
Variant classification performance of DMS-derived fitness scores and 73 missense VEPs. **(a-c)** Classification performance, measured by AUROC, for different subsets of pathogenic versus benign *ERCC2* variants, evaluated across 73 VEPs and DMS. Panel **(a)** for XP/TTD; panel **(b)** for xeroderma pigmentosum (XP); and panel **(c)** for trichothiodystrophy (TTD). The set of benign *ERCC2* variants was identical across all analyses.

## Supplemental Tables

**Table S1:** Clinically reported benign and pathogenic *ERCC2* variants, clinically reported *ERCC2* VUS, and population *ERCC2* variants from gnomAD v4.1.

**Table S2:** *ERCC2* missense variants with measured fitness scores and ACMG evidence codes assigned using reference datasets via the *acmgscaler* tool.

